# Proinsulin degradation and presentation of a proinsulin B-chain autoantigen involves ER-associated protein degradation (ERAD)-enzyme UBE2G2

**DOI:** 10.1101/2023.06.16.545333

**Authors:** Tom Cremer, Hanneke Hoelen, Michael L. van de Weijer, George M Janssen, Ana. I. Costa, Peter A. van Veelen, Robert Jan Lebbink, Emmanuel J.H.J. Wiertz

## Abstract

Type 1 diabetes (T1D) is characterized by HLA class I-mediated presentation of autoantigens on the surface of pancreatic β-cells. Recognition of these autoantigens by CD8^+^ T cells results in the destruction of pancreatic β-cells and, consequently, insulin deficiency. Most epitopes presented at the surface of β-cells derive from the insulin precursor molecule proinsulin. The intracellular processing pathway(s) involved in the generation of these peptides are poorly defined. In this study, we show that a proinsulin B-chain antigen (PPI_B5-14_) originates from proinsulin molecules that are processed by ER-associated protein degradation (ERAD) and thus originate from ER-resident proteins. Furthermore, screening genes encoding for E2 ubiquitin conjugating enzymes, we identified UBE2G2 to be involved in proinsulin degradation. These results indicate that insulin-derived peptides, presented by HLA-class I molecules at the cell surface, originate from ER-resident proinsulin that has been dislocated to the cytosol for subsequent degradation. These insights into the pathway involved in the generation of insulin-derived peptides emphasize the importance of proinsulin processing in the ER to T1D pathogenesis and identify novel targets for future therapies that may cure or even prevent T1D.

**Graphical abstract:** 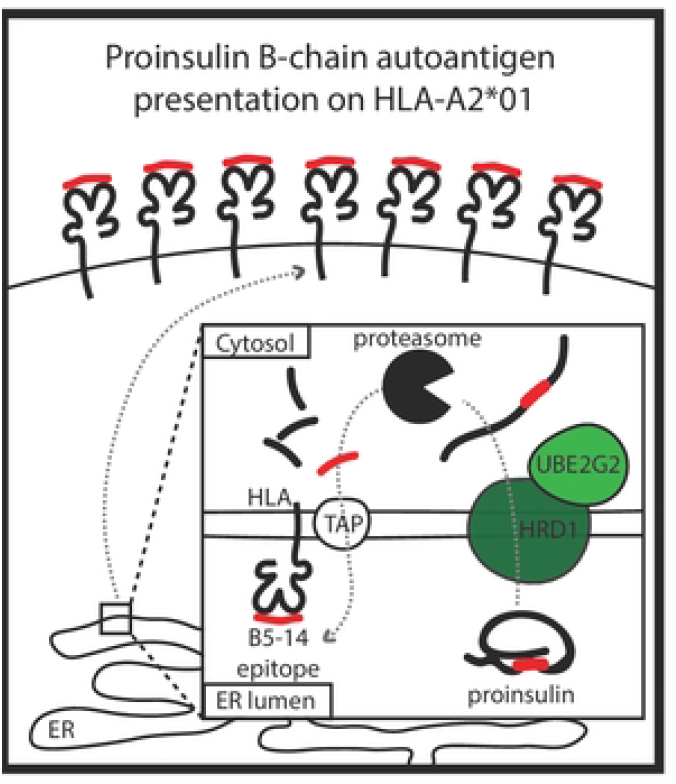

## INTRODUCTION

Presentation of autoantigens in the context of HLA class I molecules on pancreatic β-cells plays an important role in Type 1 Diabetes (T1D) pathogenesis. These autoantigens are recognized by CD8^+^ T cells, present in pancreatic islands of T1D patients [1]. Autoreactive T cells cause the destruction of a significant part of the insulin producing β-cell population, which results in severe insulin deficiency (reviewed in [2]). HLA-A^*^02:01-restricted CD8^+^ T cells have been implicated in the formation of pre-diabetic β-lesions. These T cells also accelerate the onset of T1D in HLA-A^*^02:01-transgenic NOD mice. Furthermore, the presence of this T cell population is associated with development of T1D in high-risk class II HLA-DR3 and HLA-DR4 carriers [3, 4]. The majority of epitopes recognized by autoreactive CD8^+^ T cells originate from insulin [5-7]. Evading immune responses evoked by proinsulin antigens by deleting insulin genes has been shown to prevent diabetes in the NOD mouse, emphasizing the importance of this T cell subset as a potential therapeutic target [6, 8].

Proinsulin molecules are both co- and post-translationally translocated into the ER, where they subsequently mature, starting with correct pairing of six cysteine residues to form three evolutionarily conserved disulfide bonds [9-11]. The majority of proinsulin molecules oligomerize and pass the Golgi to be sorted into secretory granules, where following C-peptide cleavage they achieve their mature form and await secretion upon glucose stimulation [10]. At the same time, terminally misfolded proinsulin molecules are removed from the ER by a quality control mechanism known as ER-associated protein degradation (ERAD) [12-14]. Erroneous proinsulin molecules are (partially) unfolded or disaggregated and prepared for ERAD by ER-resident reductases and chaperones including Grp170, PDIA, PDIA6, Calnexin and Calreticulin [13, 15-17]. Over one hundred ERAD components have been identified so far, and their complex cooperation is mostly centered around membrane-bound E3 ubiquitin ligases, which ubiquitinate ERAD substrates during dislocation across the ER membrane into the cytosol (reviewed in [18]). Upon dislocation of glycosylated substrates, asparagine-bound glycans are removed by the cytosolic enzyme *N-*glycanase [19] (illustrated in Fig. 1A), resulting in deamidation of an asparagine (N) residue to aspartate (D) (Fig. 1B). The process of attaching ubiquitin to a protein involves three steps, the first of which is activation of ubiquitin by the E1 enzyme in preparation of ubiquitin for further transfer. Subsequently, ubiquitin conjugating enzymes, or E2s, can receive activated ubiquitin from the E1. Finally, ubiquitin ligases, or E3s, select substrates for ubiquitination and facilitate transfer of one or more ubiquitin molecules to an acceptor residue [18].

**Fig. 1.**
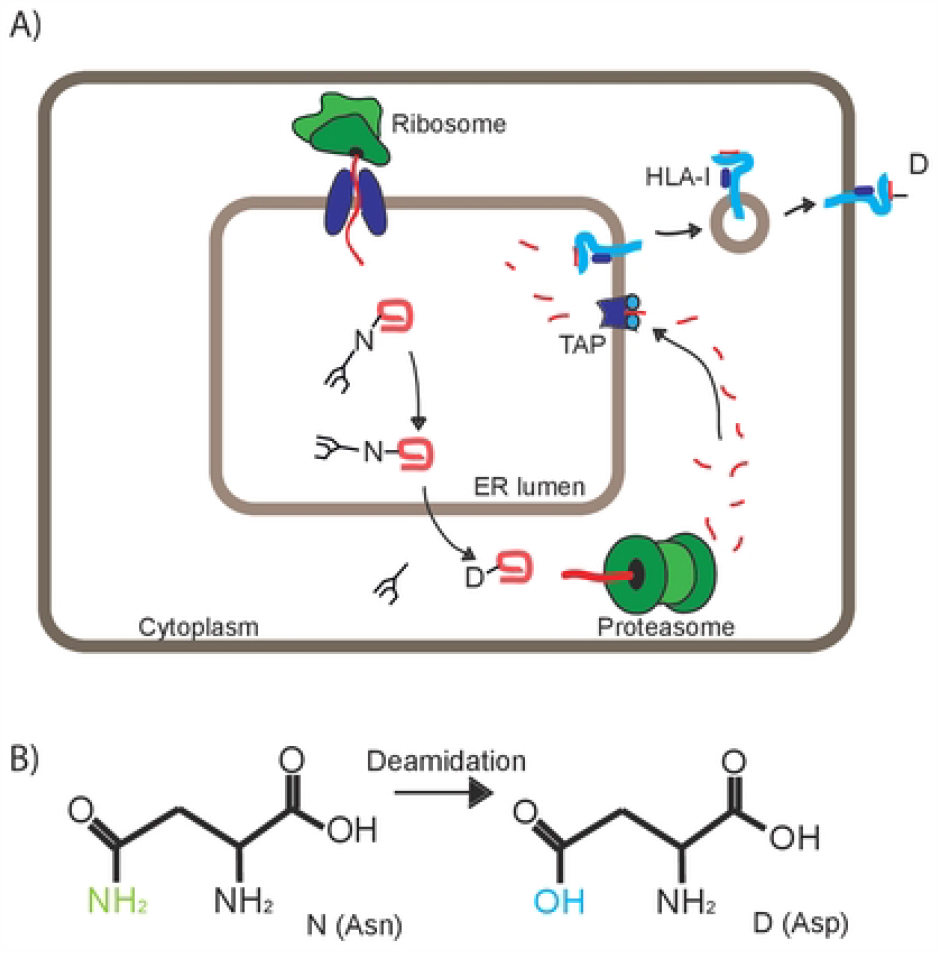
Dislocation of ERAD substrates over the ER membrane is followed by deamidation. **A**. In this study, we introduce a glycosylation consensus sequence in proinsulin to follow epitope trafficking. Schematic representation of the degradation route of PI-C31N. PI-C31N molecules are co-translationally translocated into the ER where N-linked glycosylation takes place. Dislocation of Δss-PPI-C31N molecules is accompanied by the removal of the N-linked glycan, leaving the protein deamidated. Dislocated insulin molecules are then targeted to the proteasome and the resulting proinsulin epitopes are imported into the ER by TAP, loaded onto HLA class I molecules and presented at the cell surface. **B**. Deamidation of an asparagine residue results in a aspartate residue.

More than a dozen ER-specific E3 enzymes have been identified thus far. Among these, HRD1 was found to be specifically involved in degradation of misfolded proinsulin, operating in an ERAD complex that contains Derlin-2, SEL1L and the AAA-ATPase p97 [12, 13]. P97 extracts ubiquitinated ERAD substrates from the ER membrane and shuttles them to the proteasome for degradation. In the event that proteolysis of proteasomal substrates is incomplete, this process liberates short peptides suitable for TAP-mediated re-import into the ER. Here, peptides may be loaded onto HLA class I molecules and subsequently presented to CD8^+^ T cells (Fig. 1A). Several E2 conjugating enzymes have been implicated in ERAD, including UBE2G2, UBE2J1 and UBE2J2, with the former two reported to work in conjunction with HRD1 [20-22]. The identity of the E2 enzyme that is involved in degradation of proinsulin has remained elusive so far.

In this study, we show that a B-chain proinsulin antigen presented by HLA-A^*^02:01 derives from ER-resident proinsulin molecules dislocated to the cytosol during ERAD. Additionally, we use CRISPR/Cas9-mediated knockout of HRD1 to demonstrate its involvement in degradation of proinsulin and identify UBE2G2 as an E2 conjugating enzyme for ERAD-mediated proinsulin quality control. These results shed new light on the route of proinsulin antigen processing and implicate ERAD’s molecular machinery in this process.

Identification of new therapeutic targets is of particular interest for the development of new treatment strategies and the ubiquitin conjugating and ligating enzymes identified here may serve as new strongholds in combatting (the onset of) T1D in the future.

## MATERIALS & METHODS

### Antibodies and Chemicals

Immunoprecipitation of PI was performed with a polyclonal guinea pig antibody (Millipore, #3440). Western blot analysis was performed with the following antibodies: H86 for proinsulin (Santa Cruz, cat. # sc-9168), C4 for beta-actin (Millipore, lot. #LV1728681), H68.4 for transferrin-receptor (Invitrogen, cat. # 13-6800), mouse HRD1 (Cell Signaling, #12925), D8Z4G for UBE2g2 (Cell Signaling, #63182), and HCA2 and W6/32 against HLA. Secondary antibodies used were goat α-mouse IgG(H+L)-HRP (no. 170-6516, Bio-Rad) and goat α-rabbit IgG(H+L)-HRP (no. 4030-05, Southern Biotech). Cycloheximide was from Enzo Life Sciences.

### Constructs

PPI and its glycosylation mutants and PPI-GFP were expressed from a pSico-based lentiviral plasmid [23] containing an EF1α-promotor and Zeocin and ampicillin resistance genes. CRISPR-Cas9 plasmids and E3- and E2-enzyme specific gRNAs used in the E2 screen have been described earlier [22, 23] and gRNA sequences can be found in supplementary Table 1.

### Cell culture

K562 cells stably expressing proinsulin [12] and HEK293T cells for virus production were cultured in IMDM (Lonza). All media were supplemented with 10% FCS, 100U/mL penicillin, 100U/mL streptomycin and 2mM GlutaMAX. Cell lines were all maintained in humidified incubators at 37°C and 5% CO_2_.

### Lentiviral transductions

Replication-deficient recombinant lentiviruses were produced via Mirus-mediated cotransfection of HEK293T cells with a pSico-based lentiviral vectors encoding the gene of interest, together with pCMV-VSVG, pMDLg-RRE, and pRSV-REV [12]. After 48-52 hours, supernatant was harvested. Transductions were performed by 90 minute spinoculation at 2000 RPM. Cells were allowed to recover for 48 hours after which they were selected with the appropriate antibiotic or sorted for fluorescent protein expression by FACS.

### Generation of clonal knockout cell lines with CRISPR/Cas9-mediated genome editing

K562 cells expressing HLA-A^*^02:01 and PPI were transduced with a lentiviral CRISPR/Cas9 vector, in which a single lentiviral plasmid encodes Cas9, puroR and a gRNA sequence (HRD1-1: GTGATGGGCAAGGTGTTCTT; HRD1-2; GCGGCCAGCCTGGCGCTGAC; HRD1-3: GGCCAGGGCAATGTTCCGCA; UBE2G2-1: GGAGAAGATCCTGCTGTCGG ; UBE2G2-2: GGACTTAACGGGTAATCAAG; UBE2G2-3: GACTTAACGGGTAATCAAGT). After puromycin selection (0,2mg/mL), single clones were created by limiting dilution and knock-out status was confirmed by Western blot and genomic target site sequencing. Rescue cell lines were obtained by transduction with guide-resistant cDNA expressed from a lentiviral plasmid which also encodes an mAmetrin cassette, followed by sorting of mAmetrin positive cells with a FACSAriaII (BD).

### Flow cytometry

Samples were fixed in 0,1% PFA before flow cytometric analysis by a FACSCantoII cytometer (BD). eGFP signal was quantified to assess total PPI-GFP levels. Data analysis was performed using FlowJo software (TreeStar), Excel (Microsoft), and Graphpad PRISM 6 software (Graphpad).

### Western blot

1% Triton cell lysates were treated with 50mM DTT and 100mM NEM before loading on a 12% Nu-PAGE gel (Invitrogen). Separated proteins were blotted on PVDF membranes (Millipore), blocked in 5% ELK in TBS (150mM NaCl, 3μM KCl, 25 mM Tris-HCl pH 7.5) + 5% ELK and stained with primary and secondary antibodies in TBS + 0,1% Tween-20. Detection was performed using ECL substrate solutions (Pierce) and a Imagequant LAS4000 luminescent image analyzer (GE Healthcare). Membrane scans were analyzed using ImageJ software. Statistical significance was reported from three independent, actin-normalized experiments.

### RT-PCR

mRNA transcripts were isolated using a RNeasy kit (Qiagen) and converted to cDNA using random primers and M-MuLV RT-PCR enzyme (NEB). Proinsulin-specific primers (GTGAACCAGCACCTGTGC Fw and CGGGTCTTGGGTGTGTAGAAG Rv) were used to amplify cDNA in a standard PCR reaction, which was analyzed on a 2% agarose gel.

### Peptide elution & mass-spectrometry

Approximately 1e9 cells from PPI (or variant) expressing cells were lysed in 10 ml lysis buffer (50 mM Tris-Cl pH 8.0, 150 mM NaCl, 5 mM EDTA, 0.5% Zwittergent 3-12 (N-dodecyl-N,N-dimethyl-3-ammonio-1-propanesulfonate) and protease inhibitor (Complete, Roche Applied Science)) for 2 h at 0°C (ref A). Lysates were successively centrifuged for 10 min at 2500 × g and for 45 min at 31,000 x g to remove nuclei and other insoluble material, respectively. Lysates were passed through a 100 μl CL-4B Sepharose column to preclear the lysate and subsequently passed through a 100 μl column containing 250 μg pan class I (W6/32) IgG coupled to protein A Sepharose (ref A). The W6/32 column was subsequently washed with lysis buffer, low salt buffer (20 mM Tris-Cl pH 8.0, 120 mM NaCl), high salt buffer (20 mM Tris-Cl pH 8.0, 1 M NaCl), and finally low salt buffer. HLA α chain, β2m and peptides were eluted with 10% acetic acid, diluted with 0.1% TFA and purified by SPE (Oasis HLB, Waters) by sequential elution with 20% and 30% acetonitrile in 0.1% TFA to remove HLA protein chains.

Peptides were lyophilized, dissolved in 95/3/0.1 v/v/v water/acetonitrile/formic acid and subsequently analyzed by on-line C18 nanoHPLC MS/MS with a system consisting of an Easy nLC 1200 gradient HPLC system (Thermo, Bremen, Germany), and a LUMOS mass spectrometer (Thermo). Fractions were injected onto a homemade precolumn (100 μm × 15 mm; Reprosil-Pur C18-AQ 3 μm, Dr. Maisch, Ammerbuch, Germany) and eluted via a homemade analytical nano-HPLC column (30 cm × 50 μm; Reprosil-Pur C18-AQ 3 um). The gradient was run from 2% to 36% solvent B (20/80/0.1 water/acetonitrile/formic acid (FA) v/v) in 120 min. The nano-HPLC column was drawn to a tip of ∼5 μm and acted as the electrospray needle of the MS source. The LUMOS mass spectrometer was operated in data-dependent MS/MS mode for a cycle time of 3 seconds, with a HCD collision energy at 32 V and recording of the MS2 spectrum in the orbitrap. In the master scan (MS1) the resolution was 60,000, the scan range 400-1400, at an AGC target of 400,000 @maximum fill time of 50 ms. Dynamic exclusion after n=1 with exclusion duration of 20 s. Charge states 1-3 were included. For MS2 precursors were isolated with the quadrupole with an isolation width of 1.2 Da. First mass was set to 110 Da. The MS2 scan resolution was 30,000 with an AGC target of 50,000 @maximum fill time of 100 ms.

In a post-analysis process, raw data were first converted to peak lists using Proteome Discoverer version 2.1 (Thermo Electron), and then submitted to the Uniprot Homo sapiens database (67911 entries), using Mascot v. 2.2.04 (www.matrixscience.com) for protein identification. Mascot searches were with 10 ppm and 0.02 Da deviation for precursor and fragment mass, respectively, and no enzyme was specified. Methionine oxidation was set as a variable modification. Peptides with MASCOT scores <35 were generally discarded.

## RESULTS

### HRD1 catalytic activity is involved in proinsulin degradation

To confirm the involvement of the E3 ubiquitin ligase HRD1 in proinsulin degradation [12, 13], we completely depleted cells of HRD1 by CRISPR/Cas9-mediated genome editing. K562 cells expressing HLA-A2 and PPI were transduced with guideRNAs (gRNAs) (see Table 1) targeting the region encoding for the 5’ end of the HRD1 gene. Each of the gRNAs tested reduced HRD1 protein levels in a polyclonal cell population (Fig. S1A). Clonal cell lines were established and analyzed for Cas9-mediated indel formation. Out-of-frame indels, which result in a frame-shift that may lead to a premature stop codon, were present in three selected clonal cell lines (Fig. 2A and Fig. S1B). Loss of HRD1 expression was confirmed by Western blotting (Fig. 2B and Fig. S1C). Next, we re-transduced a selected K.O. cell line to express either wild-type HRD1 or a mutant in which the first cysteine residue within the HRD1 RING domain was changed to an alanine, rendering the mutant catalytically inactive (C1A). HRD1 or HRD1 C1A expression could be detected in these cell lines (Fig. 2B and Fig. S1C). Importantly, compared to KO cells, PI steady state levels decreased in the presence of HRD1, but not HRD1-C1A. Furthermore, proinsulin is relatively stable in HRD1 KO cells, as observed in a CHX chase assay (Fig. 2C). Reintroduction of HRD1 WT, but not HRD1-C1A, enhanced proinsulin degradation (Fig. 2C and 1D), indicating that the first catalytic cysteine residue within HRD1’s RING domain is required for proinsulin degradation. These results confirm earlier studies and indicate that HRD1 catalytic activity is required for efficient degradation of proinsulin molecules.

**Fig. 2.**
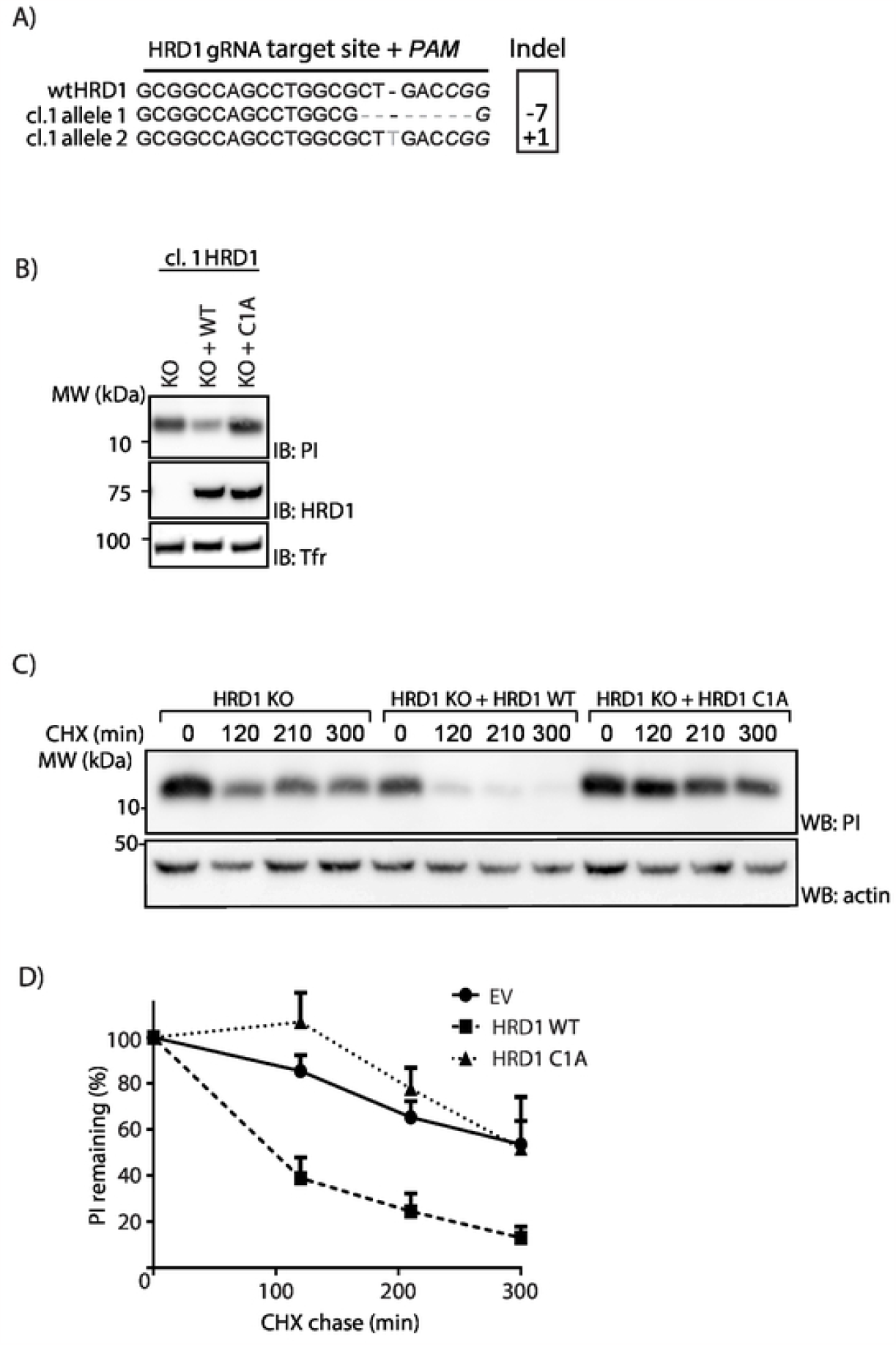
HRD1 catalytic activity is involved in proinsulin degradation. **A**. K562 cells stably expressing HLA-A^*^02:01 and PPI were transduced with a lentiviral CRISPR-Cas9 vector containing a gRNA sequence directed against the 5’ region of the HRD1 gene and selected with puromycin. Monoclonal knockouts were generated by limited dilution and analyzed for HRD1 expression. Genomic DNA was isolated and sequenced for the presence of indels within the gRNA target region. Both HRD1 alleles were aligned to a reference sequence (NCBI gene entry 84447). **B**. K562 cells from (A) were re-transduced with an empty cDNA vector (KO) or a cDNA vector encoding HRD1 (WT), or a catalytically inactive mutant (C1A). Cells were sorted on the basis of mAmetrin expression to obtain pure populations. Next, Expression of HRD1 and proinsulin was assessed in corresponding cell lysates by immunoblotting. Human transferrin receptor was used as a loading control. **C**. K562 cells from (B) were treated with 200ug/mL CHX for the indicated times, followed by WB analysis of total proinsulin levels. A representative blot from three independent experiments is shown here. **D**. Results from three independent experiments of (C) were quantified, corrected for actin levels and compared to t=0 (Error bars = SD)

### Proinsulin degradation involves the E2-ubiquitin conjugating enzyme UBE2G2

After confirming HRD1 involvement in PI degradation, we set out to identify the E2-enzyme that acts in concert with HRD1 to target proinsulin molecules to the proteasome. We used an arrayed CRISPR-Cas9 library comprised of 120 gRNAs to target 40 mammalian E2 enzymes (3 gRNAs/gene) [22]. K562 cells stably expressing a proinsulin-GFP fusion protein (PI-GFP) were transduced with E2-specific CRISPR vectors. Steady state PI-GFP levels were assessed in the gRNA-positive cell population by flow cytometry at 9 days post-infection. Comparable PI-GFP levels were detected in most knock-out cells, except in cells that expressed gRNAs targeting the ubiquitin conjugating enzyme UBE2G2 (Fig. 3A). A substantial increase of 40-50% was observed in all three cases, suggesting that UBE2G2 is an important factor in degradation of proinsulin. Notably, PI-GFP was not rescued in cells expressing gRNAs against UBE2G1, UBE2G2’s closest homologue, pointing at the specificity of the observed phenotype (Fig. 3B).

**Fig. 3.**
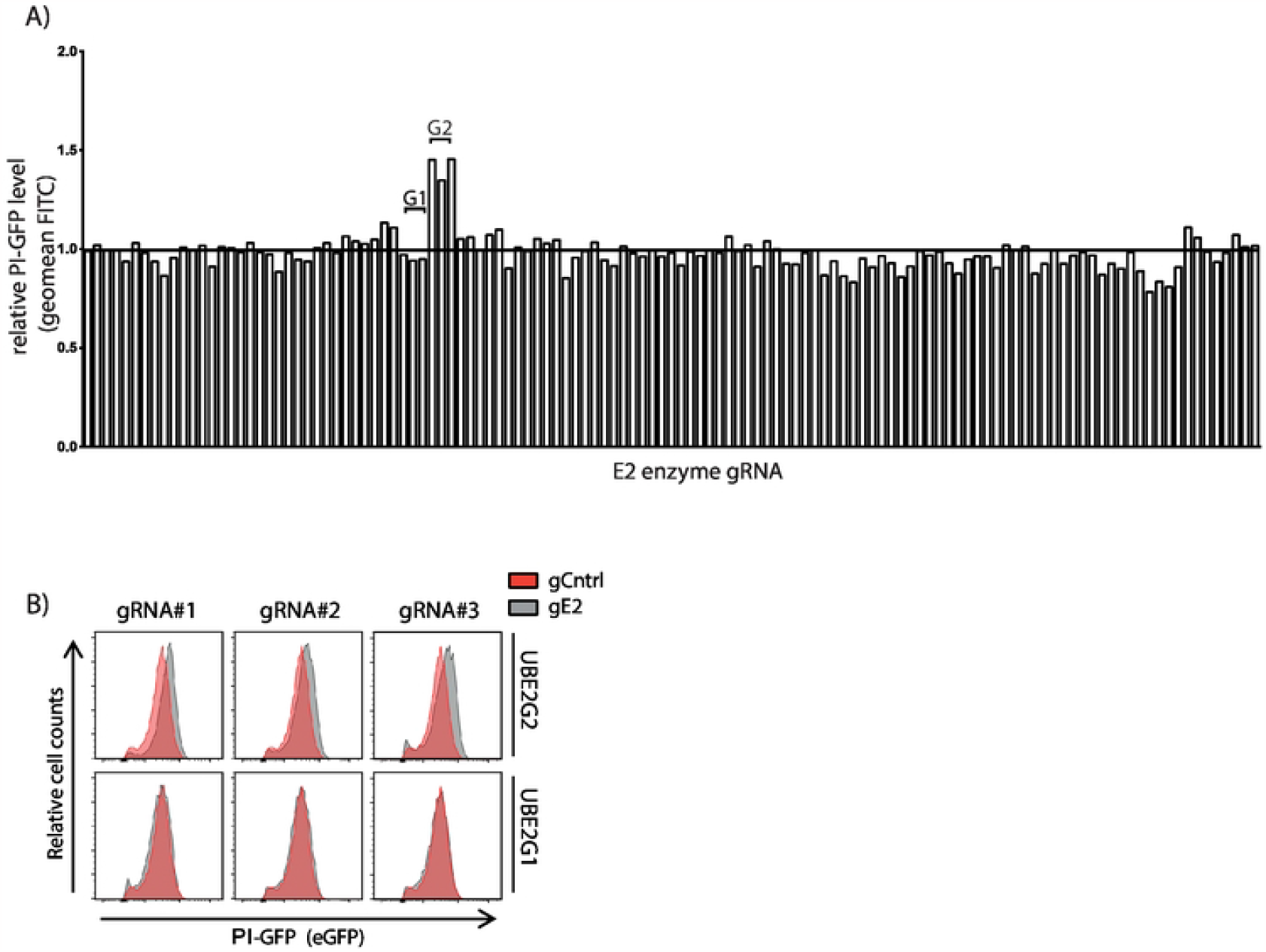
An E2-specific CRISPR knockout screen identifies UBE2G2 to be involved in PI-GFP degradation. **A**. An arrayed CRISPR library targeting every known human E2-conjugating enzyme with three different gRNAs per gene [22] was transduced into K562 cells expressing PPI-GFP. Cells were selected with puromycin and GFP levels were assessed by flow cytometry 9 days post-infection. Results were quantified and compared to GFP levels in empty vector-expressing cells. For sample IDs, see suppl. Table 1. **B**. Histograms of cells transduced with gRNAs targeting UBE2g2 or UBE2g1 (grey) or an empty vector control (red) from the screen shown in (A). PI-GFP levels were evaluated using flow cytometry.

Next, we set out to validate these results in clonal UBE2G2 knockout cells expressing untagged proinsulin. Using the gRNA that showed the most efficient depletion of UBE2G2 in the polyclonal knockout population, gRNA#1 (Fig. S2A), three clonal cell lines were created and analyzed for Cas9-mediated genome editing. We observed out-of-frame indels in selected clones (Fig. 4A and Fig. S2B), corresponding to the loss of UBE2G2 expression (Fig. 4B and Fig. S2C). Steady state proinsulin levels were increased in UBE2G2 knockout cells (Fig. 4B). To confirm that this phenotype depends on loss of UBE2G2 activity, we added back gRNA-resistant cDNAs encoding wild-type HA-tagged UBE2G2 (WT) or a catalytically inactive (C89S) mutant of UBE2G2. HA-UBE2G2 expression could be detected in these cells, and PI degradation in UBE2G2 KO cells could be restored by expression of WT UBE2G2, but not by expression of C89S UBE2G2, in a clonal setting (Fig. 4B, Fig. S2C). Importantly, as shown by a CHX chase assay (Fig. 4C, 3D), this increase in protein level is the result of a decreased proinsulin degradation rate. These results clearly indicate that UBE2G2 and its ubiquitin conjugating activity are involved in proinsulin degradation.

**Fig. 4.**
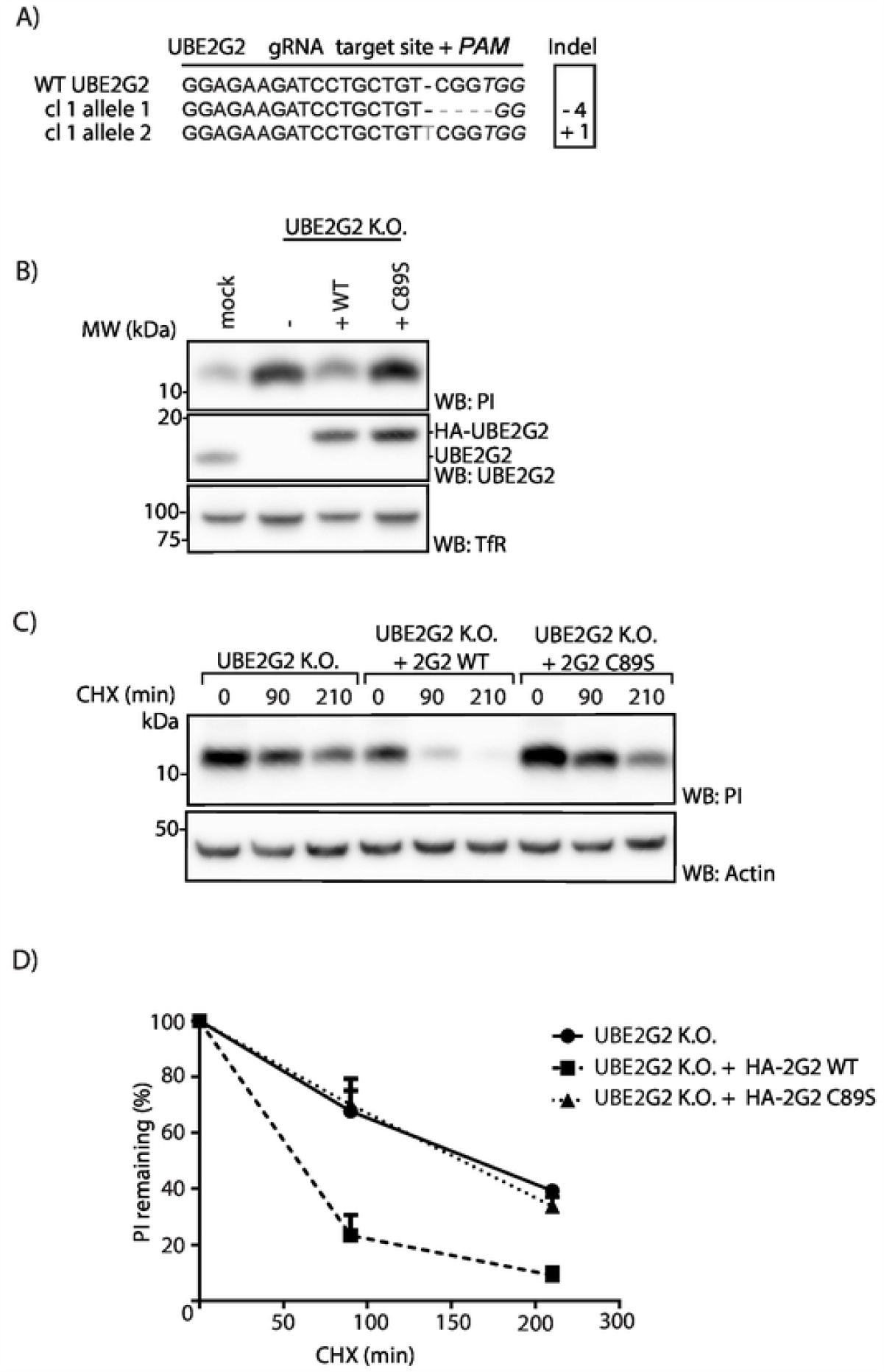
UBE2G2 catalytic activity is involved in proinsulin degradation. **A**. K562 cells stably expressing HLA-A^*^02:01 and PPI were transduced with a CRISPR-Cas9 vector containing a gRNA sequence directed against the N-terminal region of the UBE2g2 gene and selected with puromycin. Monoclonal knockouts were generated by limited dilution and analyzed for UBE2g2 expression. Genomic DNA was isolated and sequenced for the presence of indels within the gRNA target region. Both UBE2g2 alleles were aligned to a reference sequence (NCBI gene entry 7327) . **B**. K562 cells from (C) were retransduced with an empty cDNA vector (KO) or a gRNA-resistant cDNA vector encoding *wt* HA-UBE2G2 (WT), or a catalytically inactive mutant HA-UBE2G2 (C89S). Cells were sorted for mAmetrin to enrich for gRNA-positive cells. Next, cell lysates were analyzed by WB for expression of UBE2G2 and proinsulin. Human transferrin receptor (TfR) was used as a loading control. **C**. K562 cell lines from (B) were treated with 200ug/mL CHX for the indicated times, followed by WB analysis of total proinsulin levels. A representative blot from three independent experiments is shown. **D**. Results from three independent experiments of (C) were quantified, corrected for actin levels and normalized to t=0 (Error bars = SD).

### HLA-A^*^02:01 restricted presentation of the PPI_B5-14_ epitope involves dislocation of proinsulin over the ER membrane

Next, we investigated whether presentation of proinsulin-derived epitopes by HLA class I involves the ERAD machinery identified here. For this, we used a K562 cell line stably expressing HLA-A^*^02:01 [12], which presents proinsulin peptides according to the β-cell’s proinsulin-derived HLA-A2 peptidome [24]. As mentioned before, N-glycosylated ERAD substrates are deglycosylated in the cytoplasm. The intrinsic deamidation of the N-glycosylated asparagine allows us to investigate whether a proinsulin B-chain epitope (PPI_B5-14_: HLCGSHLVEA) travels from the ER back into the cytosol. To this end, we designed a preproinsulin construct which encodes an asparagine residue at position 31, thereby creating an N-glycosylated consensus motif (PPI-C31N, Fig. 5A). This motif is present in the PPI_B5-14_ peptide that is displayed on HLA-A^*^02:01 on the cell surface of our established surrogate B cell system. Involvement of ERAD in presentation of this proinsulin mutant would thus result in presentation of a peptide in which the asparagine has been converted into an aspartate residue. Three mutants were used as controls. First, a C31D mutant, which carries an aspartate residue from the beginning. Secondly, a C31N mutant lacking the signal sequence (Δss-C31N), which does not enter the ER, is not glycosylated and not subjected to dislocation, therefore subsequently leaving the asparagine residue for presentation at the cell surface. Third, a C96N mutant, which harbors a glycosylation motif within a non-presented sequence and, like C31N, lacks a cysteine residue involved in proinsulin folding. We stably expressed the different PPI constructs in K562 A2 cells (Fig. 5B). Importantly, C31N and C96N mutants were properly glycosylated, since both mutant proteins were sensitive to Endo H treatment (Fig. 5C). Notably, the glycosylated proinsulin mutants appear to be expressed at higher levels compared to wild-type proinsulin (Fig. 5B). The Δss-C31N mutant could not be detected by Western Blot or immunoprecipitation (data not shown), presumably because this cytosolic proinsulin is degraded very rapidly [25]. However, we were able to detect Δss-C31N mRNA transcripts via RT-PCR (Fig. 5D). These results indicate that all cell lines properly express and process mutant proinsulin molecules. To assess HLA content from these cells, HLA class I molecules were affinity purified and the peptide ligandome was acid-eluted and analyzed by mass-spectrometry. Peptides containing the aspartate (D) residue could be detected only on HLA molecules from cells expressing either the C31D or the C31N mutant, and not on cells expressing *wt* PI, Δss-C31N or the C96N mutant (Fig. 6A, Fig. S3). Importantly, disruption of a disulfide bond in change of a glycosylation motif did not affect presentation of PPI_B5-14_ peptides (Fig. 6A, C96N cells). The N^31^-containing peptide was predicted to bind to HLA as well as wild-type proinsulin-derived peptides (0,348 vs 0,344 for wt vs C31N, respectively, see Fig 6B) (*Net*MHC, [26]), and this was confirmed by mass-spectrometry analysis (Fig. 6A, Δss-C31N cells, Fig. S3). Altogether, these results identify ERAD as a source of the PPI_B5-14_ epitope.

**Fig. 5.**
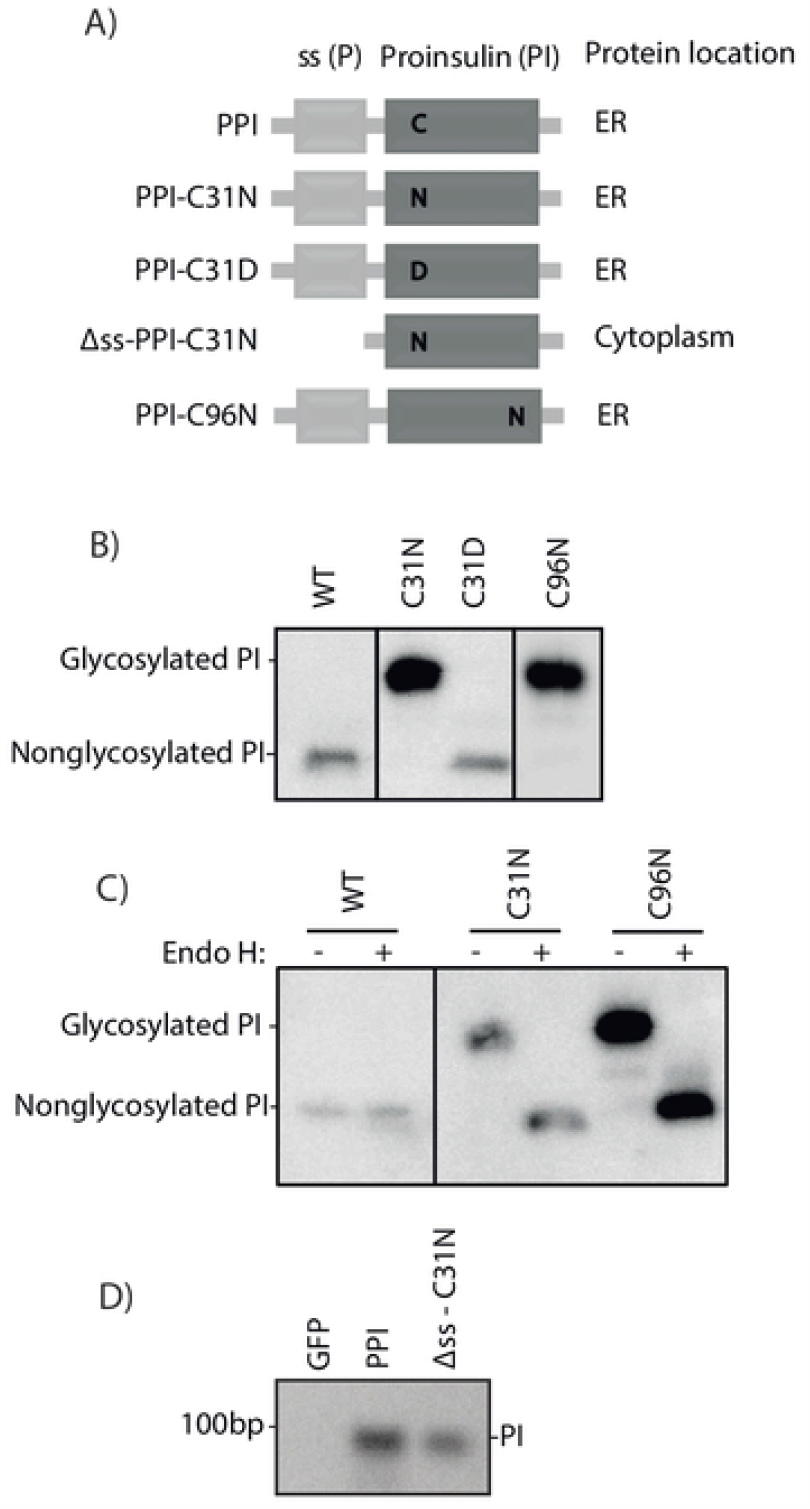
Proinsulin mutant constructs are properly expressed in K562 cells. **A**. Schematic overview of constructs that were used in the experiments in this figure and mass-spectrometry analysis in Fig. 6. **B**. Expression of mutant proinsulin constructs in cells detected by Western blot. Shorter exposure for C96N mutant is shown because of high protein levels. **C**. Cell lysates were treated with Endo H before Western blot analysis of the removal of N-linked sugar groups from glycosylation proinsulin mutants. **D**. Detection of expression of Δss-PPI-C31N mRNA species by RT-PCR.

**Fig. 6.**
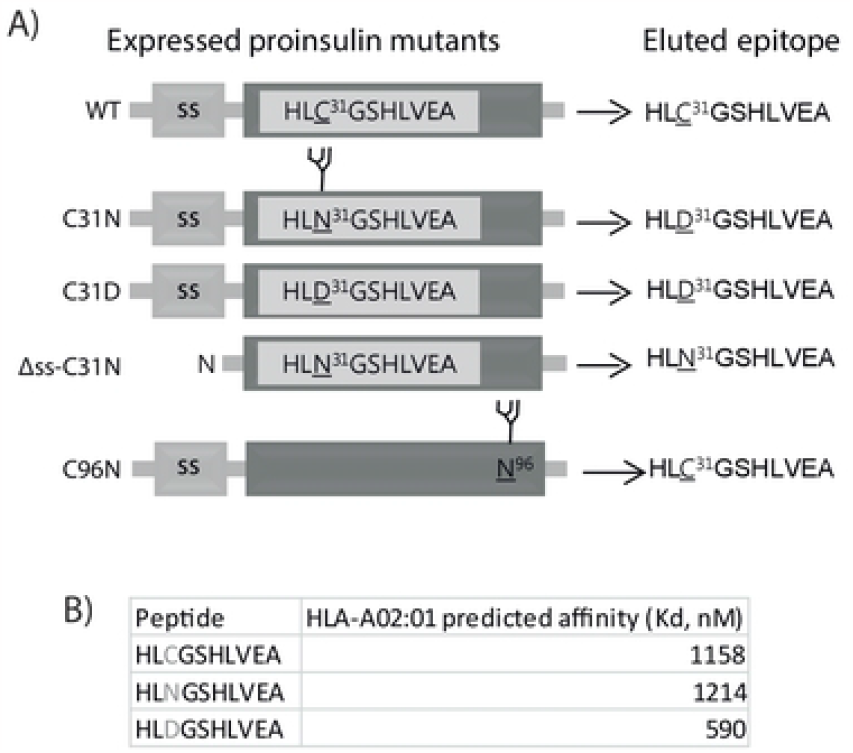
Presentation of the PPI_B5-14_ epitope requires dislocation of proinsulin over the ER membrane. **A**. Representation of mass-spectrometry results of HLA-bound peptides that were eluded from affinity-purified MHC molecules derived from K562 HLA-A2 cells expressing the indicated mutant proinsulin constructs. **B**. Affinity of peptides that result from processing of mutant proinsulin constructs for HLA-A^*^02:01 molecules as predicted with NetMHC 3.4 (http://www.cbs.dtu.dk/services/NetMHC/)

## DISCUSSION

In this study, we tracked the origin of clinically important B-chain peptides that are presented on the cell surface of pancreatic β-cells. We propose a model that involves dislocation of proinsulin from the ER into the cytosol, where it is degraded by the proteasome. In β-cells, this may occur if the ER is overloaded, as part of the unfolded protein response (UPR). β-cells produce large quantities of insulin whose biosynthesis requires cleavage of the PPI precursor and correct formation of multiple disulfide bonds. Any deviation from proper assembly results in accumulation of misfolded proinsulin, which triggers the UPR. As part of this response, ERAD-mediated dislocation of proinsulin will be initiated to relieve the ER from stress. At the heart of ERAD is ubiquitination of substrates by specific combinations of ubiquitin conjugating and ligating enzymes [18], and the work presented here indicates the involvement of the ERAD E3 enzyme HRD1 and the E2 enzyme UBE2G2 in proinsulin degradation. In the cytosol, dislocated glycoproteins lose their N-linked glycans by a cytosolic N-glycanase, which results in deamidation of the asparagine residue the glycan was attached to [19]. The proteasome degrades these substrates into peptides, which may return into the ER lumen via TAP and may subsequently be loaded on HLA class I molecules. We made N-glycosylated mutant forms of proinsulin to demonstrate that presentation of a proinsulin B-chain peptide requires dislocation of proinsulin into the cytosol. Presentation of the B-chain epitopes on HLA-A^*^02:01 and the cytotoxic function of the autoreactive CD8^+^ T cell population it evokes are of particular interest, since it has been shown that these factors play a major role during the early stages of T1D pathogenesis [3, 27].

Since ER-targeting of proinsulin was shown to drastically alter antigen presentation efficiency of proinsulin-derived peptides, identification of the intracellular origin of HLA class I-presented epitopes in β-cells is of significant importance [25]. For decades, this issue has been a subject of intense debate. Proponents of the Defective Ribosomal Product (DRiP) hypothesis (reviewed in [28]) have claimed that HLA class I epitopes are mainly derived from immature, newly synthesized proteins or defective products that may have initiated from internal ribosomal entry sites, all of which are prone for proteasomal degradation. DRiPs stemming from secretory proteins may form either as a consequence of flawed translation onset in the cytoplasm or erroneous translocation into the ER. Opponents of the DRiP theory accredit normal turnover of fully functional proteins (also called ‘retirees’) as the primary source of peptides presented by HLA class I [29]. Here, we show that a B-chain proinsulin epitope originates from dislocated proinsulin species rather than from **1)** cytosolic degradation of a previously described pool of misfolded, and thus vulnerable, cytosolic preproinsulin molecules translocated post-translationally in a Sec62-dependent fashion, **2)** proinsulin peptides that result from flawed translation onset in the cytoplasm, or **3)** a recently uncovered RTN3-dependent ERphagy pathway during which proinsulin remains unexposed to cytosolic N-glycanase [9, 17, 30]. Furthermore, since mature insulin molecules reside in β-cell secretory granules, and are therefore unlikely to be targeted to ERAD for degradation, the PPI B-chain epitope is unlikely to be a derivative of long-lived insulin molecules (i.e. insulin retirees). Thus, along the lines of the DRiP hypothesis, our data strongly suggest that proinsulin epitopes, such as PPI_B5-14_, derive from immature proinsulin molecules that may have failed to achieve a proper conformation required for ER export and are consequently subjected to dislocation via ERAD.

Previous studies on factors involved in proinsulin degradation have focused on the misfolded C96Y proinsulin mutant expressed in Akita diabetic mice. This model has been used to study Mutant Insulin gene-induced Diabetes of Youth (MIDY). Here, CRISPR knockout has been used to validate the involvement of HRD1 in wild-type proinsulin degradation. Furthermore, the E2-specific CRISPR/Cas9 screen presented here identified UBE2G2, but not other ERAD-associated E2’s UBE2J1 or UBE2J2, as an essential E2 ubiquitin conjugating enzyme responsible for proinsulin degradation. Although HRD1 has been shown to join forces with UBE2J1 [20], our data now suggest that HRD1 and UBE2G2 can also cooperate in mammalian ERAD. This is in line with a recent model in which UBE2G2 is responsible for elongation of ubiquitin chains on substrates that are primed by UBE2J1 family member UBC6, but can also independently drive poly-ubiquitination on ERAD substrates [31]. As assayed by CHX chase in this study, and by radioactive labeling earlier [12], K562 cells degrade wild-type proinsulin at a rate that is comparable to the reported degradation rate of misfolded Akita proinsulin in 293T cells and β-cells [13, 32]. Combined, these results suggest that a large portion of newly synthesized wild-type proinsulin is misfolded in K562 cells (similar to Akita disulfide mispairing), presumably because our model system lacks the β-cell-specific proinsulin folding machinery. Since β-cells synthesize enormous quantities of proinsulin molecules, it is not surprising that a substantial proportion of molecules encounters problems in attaining a mature conformation and is selected for degradation. This cascade may eventually result in presentation of proinsulin-derived autoantigens, such as PPI_B5-14_, on the β-cell surface. While the ER of K562 cells may not reflect the natural proinsulin folding environment, these cells display relevant PPI-derived epitopes on HLA-A^*^02:01 and thus resemble a genuine β-cell regarding the pathways involved in generation of these autoantigens.

In a subset of T1D patients, proinsulin has been found to accumulate in the ER [33]. Accumulation of incorrectly folded proinsulin leads to ER stress, and this has been shown to play a significant role in the production of neo-or auto-antigens, β-cell destruction and (the onset of) T1D (reviewed in [34, 35]). Correct folding of proinsulin and its exit from the ER mainly depend on formation of at least two out of three conserved disulfide bonds, and the ER of β-cells harbors specialized enzymes including PDI and ERO1β to catalyze this process [10, 34]. Modulation of the abundance of these proteins has been shown to enhance proinsulin secretion and relieve the β-cell from ER stress [15, 36]. Next to these redox enzymes, factors such as SDF2L1 [14] and Grp170 [17, 32, 37] are implicated in chaperone complexes that aid in proinsulin folding, shifting proinsulin homeostasis towards export versus accumulation and degradation. Our data strongly suggests that degradation of proinsulin through HRD1 and UBE2G2-mediated ERAD results in presentation of a proinsulin B-chain autoantigens. In this light, reducing the proinsulin degradation load on ERAD by improving the β-cell’s oxidative folding environment may decrease the number of self-peptides available in the prediabetic pancreas for presentation to autoreactive CD8^+^ T cells. Since another B-chain antigen (PPI_B10-18)_ has been implicated in graft rejection after islet transplantation [27], intervening with its cell surface presentation, for instance via genetic modification [38], may improve graft acceptance. An approach as mentioned above may concurrently lower ER stress, improve proinsulin maturation and export, and decrease ERAD-mediated presentation of proinsulin autoantigens in β-cells. Such a tactic would hit three birds with one stone in combatting T1D.

## ACKNOWLEDGEMENTS

We would like to thank dr. Ilana Berlin (LUMC) and dr. A. Zaldumbide (LUMC) for critical reading of the manuscript.

## AUTHOR CONTRIBUTIONS

TC and HH performed the experiments. GJ and PvV performed and analyzed mass spectrometry experiments. MvdW compiled the E2 gRNA library and provided technical support. TC, AC and EW wrote the manuscript. RJL and EW supervised the project. EW is the guarantor of this work and, as such, had full access to all the data in the study and takes responsibility for the integrity of the data and the accuracy of the data analysis.

## DATA AND RESOURCE AVAILABILITY

The data sets generated during the current study are available from the corresponding author upon reasonable request

